# Low back pain among nurses working in a clinical settings of Africa: A systematic review and meta-analysis of a 19 years of studies

**DOI:** 10.1101/507053

**Authors:** Ayele Semachew, Yinager Workineh, Emiru Ayalew, Worku Animaw

**Author notes:** Corresponding author Email addresses: Ayele Semachew (AS) or, Yinager Workineh (YW), Emiru Ayalew (EA), Worku Animaw (WA).

## Abstract

**Introduction:** Because of the nature of the work, healthcare providers are prone to develop different musculoskeletal problems including low back pain and hospital healthcare workers are groups of healthcare workers who suffered a lot from it. The incidence varies between countries and professions. The situation is somewhat worsen among the frontline healthcare provider in many healthcare facilities. Nurses in Africa are arguably the most important frontline health care workers available in most African healthcare facilities, performing a broad range of tasks and working in settings where no other health workers, including physicians, are available. This situation is considerably important in the causation of work load. Nursing is listed among the highly risky profession for the development of low back pain and has been ranked with in the top tenth professions which have a great risk of having susceptible to low back pain.

**Objective:** The aim of this systematic review and meta-analysis was to ascertain whether LBP is of a significant concern among nurses in African healthcare facilities.

**Methods:** A comprehensive literature search of different data bases with no date limit was conducted from September to November 2018 using the PRISMA guideline. The quality of the included studies were assessed using a 12-item rating system. Subgroup and sensitivity analysis were performed. Cochran’s Q and the I^2^ test were used to assess heterogeneity. The presence of publication bias was evaluated by using Egger’s test and visual inspection of the symmetry in funnel plots.

**Result:** During the period 2000–2018, nineteen studies with a sample size of 6110 have been carried out. Among them, the lowest and the highest prevalence were found to be 44.1% and 82.7%. Both the highest and the lowest prevalence of low back pain were reported from a studies done in Nigeria. The estimation of the prevalence rate of low back pain among nurses using the random effects model was found to be 64.07% (95% CI: 58.68–69.46; P-value < 0.0001). Heterogeneity of the reviewed studies was I^2^ = 94.2% and heterogeneity Chi-squared = 310.06 (d.f = 18), P-value < 0.0001. The subgroup analyses showed that the highest prevalence of LBP among nurses was from west African region with prevalence rates of 68.46% (95% CI: 54.94–81.97; P-value <0.0001) and followed by north Africa region with prevalence rate of 67.95% (95% CI: 55.96–79.94; P value <0.0001) had the higher prevalence of LBP as compared to their south African counterparts, 59% (95% CI: 51–66.9; P-value <0.0001).

**Conclusion:** Even though the overall prevalence of the present study is lower when compared to the western and Asian studies, it indicated that the prevalence of low back pain among nurses is on the move.

## Introduction

Low back pain (LBP) is one of the most common causes of musculoskeletal disorders related to work status (1). LBP is a rampant health problem responsible for serious suffering and disability than any other health complaint across the globe (2). It is estimated that LBP may be experienced as much as 80% in different population groups at some time in their lives (3–7). LBP has been shown to account for an average number of disability-adjusted life years (DALYs) higher than different infectious diseases, non-communicable disease and road traffic injuries. According to the Global Burden of Disease (GBD) 2010 report, LBP was recorded among the top ten high burden diseases and injuries (8).

Because of the nature of the work, healthcare providers are prone to develop different musculoskeletal problems including LBP (9)(10)(11) and hospital healthcare workers are groups of healthcare workers who suffered a lot from it. The incidence varies between countries and professions (12,13). The situation is somewhat worsen among the frontline healthcare provider in many healthcare facilities (14).

Nurses in Africa are arguably the most important frontline health care workers available in most African healthcare facilities, performing a broad range of tasks and working in settings where no other health workers, including physicians, are available (15). This situation is considerably important in the causation of work load. Nursing is listed among the highly risky profession for the development of LBP and has been ranked with in the top tenth professions which have a great risk of having susceptible to LBP (16–19). A study done in American indicated that nurses are ranked the sixth highest with regard to lose their working days from job due to LBP (20).

In providing a care, nurses are subjected to lift and transport patients or equipment, often in difficult environment particularly in developing nations where lifting aids are not available or practicable (21–25). This process is really hard on a nurse’s back(26). Biomechanical investigations reported that such movements result into high spinal stresses (27).

Lower back pain directly affects nurses’ productivity at work and consequently reduces the overall amount and quality of healthcare the clients receive (28–33). In addition, LBP will have many negative impact on different aspects of the healthcare system including absence from work place, loss of optimal performance, low job satisfaction, rising medical costs and occupational disability (34). The health of nurses influences not only their job satisfaction, quality of life and desire to change careers but also quality of care and patient safety (9,35–37).

LBP has been identified as one of the main causes of loss of hours and days among the working class citizens (38). Describing the extent of musculoskeletal injury in nurses, survey showed that nurses lost 750,000 days a year as a result of back pain (24).

Different epidemiological studies have been done to identify and relate possible risk factors to the occurrence of LBP among nursing staffs. They found that individual factors such as age, gender, educational level, body mass index, and psychosocial factors referring to job satisfaction, work stress, and anger have been examined and related to the occurrence of LBP. But, LBP is a complex condition with several factors contributing to its occurrence (17,39–44). Psychosocial factors (low work support from superiors and poor nurse physician communication) are stated as an important underlying factors for the development of LBP (21,45–47).

To the researchers’ knowledge, there is no prior systematic review and meta-analysis reporting on the prevalence of LBP among nurses. Hence, the objective of the current review was to thoroughly evaluate peer-reviewed published studies on the reported prevalence of LBP among nurses working at different African healthcare facilities. This would help us to ascertain whether LBP is of a significant concern among nurses in African healthcare facilities. In addition, the review tried to assess the methodological relevance of the retrieved studies in the subject area and this would help in identifying chances for service improvement in the African healthcare settings.

## Method

This systematic review and meta-analysis was conducted using studies that addressed low back pain among nurses working at different African healthcare facilities and the review was presented using the PRISMA guideline (48).

### Search Strategy

A comprehensive search was conducted from September to November 2018 with no date limit to each data bases. Electronic searches using main sets of terms and using their combinations was performed. The first sets were the key words that describes the population under study, the next sets of terms includes the outcome of interest of the study, the other sets of terms were the settings of the study and the final sets of terms were the study area. Based on this principle, electronic data base searching was done. The search was done with the phrase/Boolean search mood from the title, abstract and keywords. Searching terms and their combinations were; (“Nursing” OR “Professional Nurses” or “Nurses”) AND (“Low back pain” OR “Prevalence of low back pain” OR “Incidence of low back pain” OR “Musculoskeletal problem”) AND “Hospitals” OR “Healthcare facilities” AND “Africa” OR “African countries”

The following data bases: PubMed, ScienceDirect, Google scholar, MEDLINE, CINHAL and ProQuest were searched to search the eligible studies. In addition to the electronic database searches, a secondary search technique known as “footnote chasing” was utilized to identify additional articles for inclusion in the review.

### Eligibility criteria

Primary research works that reported the prevalence of LBP among nurses working at different African healthcare facilities were included. Studies were not restricted by time of study or year of publication but they should be written or published using English language. Thesis report, dissertation, and any report proceedings/conference in the subject matter which was published in journals were included in our searches.

### Definitions of terms

Musculoskeletal disorders: Any pain or discomfort in one or more limbs. Low Back Pain (LBP): Any pain in the lower back between L1 - L5 (lumbar spine) and L5-S1 (lumbosacral joint) (49).

Prevalence of low back pain: A 12-month recall period was used for experiencing of low back pain, as this has been shown to be an appropriate time-scale in other studies (50).

Nurses: Nurses working at different hospitals of African countries.

### Data extraction

To extract the data, a form was prepared that included the following variables: Author names, year of publication, country, region in the continent, setting, study design, sample size, gender, mean age, measurement, prevalence of LBP, and studies’ quality score. The extraction was done by three independent researchers (AS, YW and EA). When there was disagreement between them, a thorough discussion was made between them and if there was still any disagreement, the fourth author (WA) was consulted.

### Study quality assessment

To assess the quality of the included studies, a modified critical appraisal tool was utilized. This tool includes three methodological tests containing 12 discrete criteria for prevalence studies. From these, three questions assesses sample representativeness of the target population, six questions assesses the data quality, and the remaining three questions assesses the definition of the outcome variable. Based on this parameter, studies with at least 75% of the total score were acceptable (49,51–53) to be included to the systematic review and meta-analysis (Appendix).

### Statistical analysis procedure

Data analysis was performed using STATA version 11 software and P-value ≤ 0.05 significance level was considered. The weight given to each study was assigned according to the inverse of the variance. Cochrane Q and I^2^ statistics were used to assess heterogeneity among studies. Heterogeneity was measured by I^2^ and divided into four categories; no heterogeneity (0%), low (25–50%), moderate (50–75%), and high (>75%) (54).

Subgroup analysis and meta-regression (the relationship between the years of the study and region in the continent with the prevalence rate) were employed to explore the cause of heterogeneity between studies. Funnel plot (Begg’s test) and Egger’s statistics with pseudo 95% confidence interval was used to examine publication bias.

## Result

Until December 10, 2018 four hundred eighteen articles were identified, all records were reviewed and 361 irrelevant and duplicate studies were excluded. The full texts of the remaining 57 articles were reviewed in detail and finally 19 articles met the inclusion criteria and included to the final section of the analysis (Figure 1).

**Figure 1:**
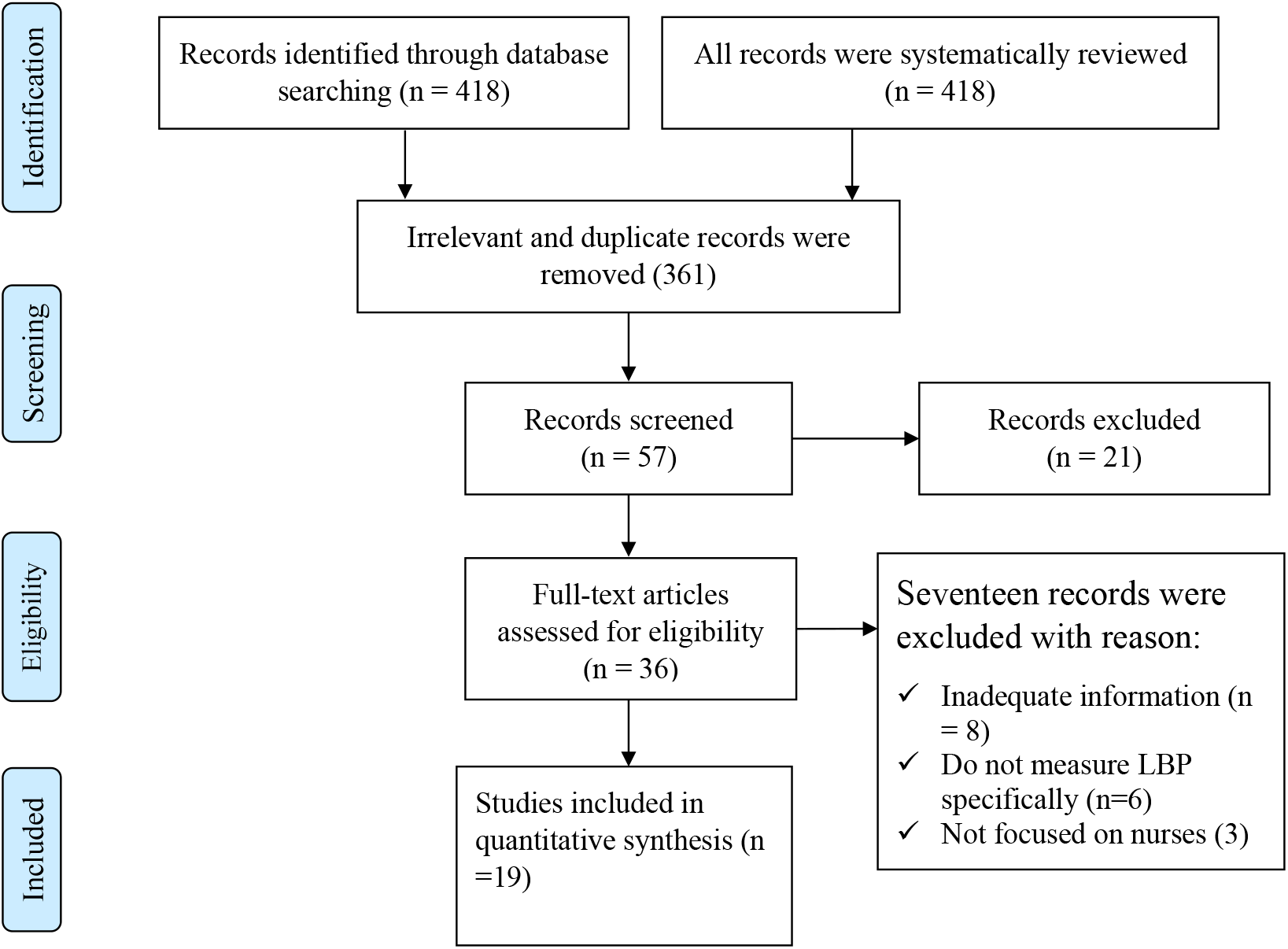
PRISMA flow diagram on prevalence of Low Back Pain (LBP) among nurses working at different healthcare facilities in Africa, 2018.

### Critical appraisal result of the included studies

A valid and standardized questionnaires were the main data collection tools in the included studies because of this, criterion number 8 and 9 in the selected critical appraisal instrument were not applicable for most studies except studies done by (55) as they utilized both interview and physical examination techniques to gather the data. Studies done by (31)(56)(57) utilized both self-administered questionnaire and physical examination so that they utilized criterion number 9 as a critical appraisal item. All the included studies for this systematic review and meta-analysis were methodologically assessed and they satisfied the indicated criteria (Table 1).

**Table 1:** Critical appraisal result of the included studies

| Included articles | Criterion No. |  |  |  |  |  |  |  |  |  |  |  |  |
| --- | --- | --- | --- | --- | --- | --- | --- | --- | --- | --- | --- | --- | --- |
|  | 1 | 2 | 3 | 4 | 5 | 6 | 7 | 8 | 9 | 10 | 11 | 12 | % |
| Thembelihle D. et al.(58) | ✓ | ✓ | ✓ | ✓ | ✓ | ✓ | ✓ | NA | NA | X | ✓ | ✓ | 90 |
| Asmare Y. et al.(55) | ✓ | X | ✓ | ✓ | ✓ | ✓ | ✓ | ✓ | ✓ | X | ✓ | ✓ | 83 |
| M M. Belay et al.(59) | ✓ | ✓ | ✓ | ✓ | ✓ | ✓ | ✓ | NA | NA | X | ✓ | ✓ | 90 |
| Lamina S. et al.(21) | ✓ | X | ✓ | ✓ | ✓ | ✓ | ✓ | NA | NA | X | ✓ | ✓ | 80 |
| Lamina S. et al.(21) | ✓ | X | ✓ | ✓ | ✓ | ✓ | ✓ | NA | NA | X | ✓ | ✓ | 80 |
| F. O. Omokhodion et al.(60) | ✓ | ✓ | ✓ | ✓ | ✓ | ✓ | ✓ | NA | NA | X | ✓ | ✓ | 90 |
| Sikiru L & Hanifa S(61) | ✓ | ✓ | ✓ | ✓ | ✓ | ✓ | ✓ | NA | NA | ✓ | ✓ | ✓ | 100 |
| Muhammed A. et al(28) | ✓ | X | ✓ | ✓ | ✓ | ✓ | ✓ | NA | NA | X | ✓ | ✓ | 80 |
| Mukaruzima Lela(62) | ✓ | ✓ | ✓ | ✓ | ✓ | ✓ | ✓ | NA | NA | X | ✓ | ✓ | 90 |
| Thembelihle D.(63) | ✓ | X | ✓ | ✓ | ✓ | ✓ | ✓ | NA | NA | X | ✓ | ✓ | 80 |
| Chandeu Mwilila(64) | ✓ | ✓ | ✓ | ✓ | ✓ | ✓ | ✓ | NA | NA | ✓ | ✓ | ✓ | 100 |
| Wided B. et al(31) | ✓ | X | ✓ | ✓ | ✓ | ✓ | ✓ | NA | ✓ | X | ✓ | ✓ | 82 |
| Ian G Munabi. et al.(65) | ✓ | X | ✓ | ✓ | ✓ | ✓ | ✓ | NA | NA | X | ✓ | ✓ | 80 |
| Mengestie M. et al.(59) | ✓ | X | ✓ | ✓ | ✓ | ✓ | ✓ | NA | NA | X | ✓ | ✓ | 80 |
| Betty C.(26) | ✓ | X | ✓ | ✓ | ✓ | ✓ | ✓ | NA | NA | X | ✓ | ✓ | 80 |
| Amany M. et al.(56) | X | X | ✓ | ✓ | ✓ | ✓ | ✓ | NA | ✓ | ✓ | ✓ | ✓ | 82 |
| Ziadi B. et al.(66) | ✓ | X | ✓ | ✓ | ✓ | ✓ | ✓ | NA | NA | X | ✓ | ✓ | 80 |
| Bolanle MS. et al.(57) | ✓ | ✓ | ✓ | ✓ | ✓ | ✓ | ✓ | NA | ✓ | X | ✓ | ✓ | 91 |
| Chiwaridzo et al.(67) | ✓ | X | ✓ | ✓ | ✓ | ✓ | ✓ | NA | NA | X | ✓ | ✓ | 80 |
| v= criterion fulfilled, X = criterion not fulfilled NA = not applicable |  |  |  |  |  |  |  |  |  |  |  |  |  |

Based on the exclusion and inclusion criteria, 19 studies ((26,28,31,38,44,55,58,61,60,62–65,59,56,66,57,67)) were included in the final analysis. All studies were done using a cross sectional study design and within the included 19 studies, a total of 6110 nurses participated. Regarding the study participants, most of them were females even if some studies (21,28,31,66) failed to report number of male and female participants in their studies clearly. Whereas a study done by (56) only incorporated female participants in their study. The sample size of the studies ranged between 80 which was a study done in Nigeria (60) and 880 a study conducted in Uganda (65). Concerning to the study settings, almost all were conducted in a hospital basis except one study which was done in Ethiopia, in addition to hospital nurses investigators incorporated nurses from health centers (55) (Table 2).

**Table 2:** Characteristics of included articles in the systematic review and meta-analysis, 2018.

| Authors name | Year of publication | Country | Region in the continent | Setting | Study design | Sample size | Sampling method | Measurement | Gender (%) | Mean age | Response rate (%) | No. of people with the outcome | Prevalence (%) |
| --- | --- | --- | --- | --- | --- | --- | --- | --- | --- | --- | --- | --- | --- |
| Thembelihle D. et al | 2018 | South Africa | South | Regional Hospital | cross-sectional | 373 | Convenience | SSAQ | 89% Female | ND | 65.0 | 157 | 59 |
| Asmare Y. et al. | 2015 | Ethiopia | East | Public Hospital & HC | cross-sectional | 428 | Survey | A-NMSQ | 53.7% Female | 30 | 91% | 222 | 57.1 |
| M M. Belay et al. | 2016 | Ethiopia | East | Public Hospital | cross-sectional | 430 | SRS | SSAQ + VAS | 72.2% Female | 30.6 | 92% | 181 | 45.8 |
| Lamina S. et al. | 2009 | Ethiopia | East | Public Hospital | cross-sectional | 120 | Convenience | SSAQ | ND | ND | 83% | 60 | 60 |
| Lamina S. et al. | 2009 | Nigeria | West | Specialized hospital | cross-sectional | 500 | Convenience | SSAQ | ND | ND | 82% | 300 | 73.5 |
| F. O. Omokhodion et al. | 2000 | Nigeria | West | Rural hospital | cross-sectional | 80 | Convenience | SSAQ | 33.8% Female | 43.8 | 93% | 51 | 69 |
| Sikiru L & Hanifa S | 2010 | Nigeria | West | Specialized hospital | cross-sectional | 500 | ND | SSAQ | 63.7% Female | 39.2 | 82% | 300 | 73.5 |
| Muhammed A. et al | 2015 | Nigeria | West | UTH | cross-sectional | 120 | multi-stage | SSAQ | ND | ND | 82% | 81 | 82.7 |
| Mukaruzima Lela | 2010 | Rwanda | East | Military Hospital | cross-sectional | 133 | SRS | IPAQ+ NMDQ | 82% Female | 34.5 | 82% | 88 | 78 |
| Thembelihle D. | 2010 | South Africa | South | Public Hospital | cross sectional | 373 | SRS | SSAQ | 79.3%Female | ND | 72% | 158 | 59 |
| Chandeu Mwilila | 2008 | Tanzania | East | MOI | cross sectional | 312 | Purposive | SSAQ | 83.6%Female | 35.9 | 54% | 126 | 73.6 |
| Wided B. et al | 2017 | Tunisia | North | Teaching hospital | cross-sectional | 329 | ND | Borg CR-10 scale, JCQ | ND | 39.8 | 61.70% | 125 | 58.1 |
| Ian G Munabi. et al. | 2014 | Uganda | East | Public Hospital | cross-sectional | 880 | ND | DMQ & NMDQ | 85.7%Female | 35.4 | 85.40% | 459 | 61.9 |
| Mengestie M. et al. | 2016 | Ethiopia | East | Public Hospital | cross-sectional | 395 | SRS | SSAQ | 72.2%Female | 30.6 | 91.9 | 166 | 45.8 |
| Betty C. | 2015 | Kenya | East | Both public & private hospitals | cross-sectional | 169 | SRS | SSAQ | 76.9%Female | 35 | 76.9 | 100 | 76.9 |
| Amany M. et al. | 2014 | Egypt | North | Public Hospital | cross-sectional | 150 | Purposive | OLBDQ | 100%Female | ND | 100 | 119 | 79.3 |
| Ziadi B. et al. | 2014 | Algeria | North | Public Hospital | cross-sectional | 450 | SRS | SSAQ | ND | ND | 66.7 | 200 | 66.7 |
| Bolanle MS. et al. | 2010 | Nigeria | West | UH,GH,PH | cross-sectional | 160 | SRS | SSAQ | 97.5%Female | 36.4 | 80 | 56 | 44.1 |
| Chiwaridzo et al. | 2018 | Zimbabwe | East | Public Hospital | cross-sectional | 208 | Stratified RS | NMDQ | 84.6%Female | 32 | 55.7 | 65 | 55.7 |

### Prevalence of low back pain (LBP)

During the period 2000–2018, nineteen studies with a sample size of 6110 have been carried out. Among them, the lowest and the highest prevalence were found to be 44.1% and 82.7%. Both the highest and the lowest prevalence of LBP were reported from a studies done in Nigeria. The lowest prevalence of LBP was reported by (57) whereas the highest prevalence was reported by (28). The estimation of the prevalence rate of LBP among nurses using the random effects model was found to be 64.07% (95% CI: 58.68–69.46; P-value < 0.0001). Heterogeneity of the reviewed studies was I^2^ = 94.2% and heterogeneity Chi-squared = 310.06 (d.f = 18), P-value < 0.0001 (Figure 2).

**Figure 2:**
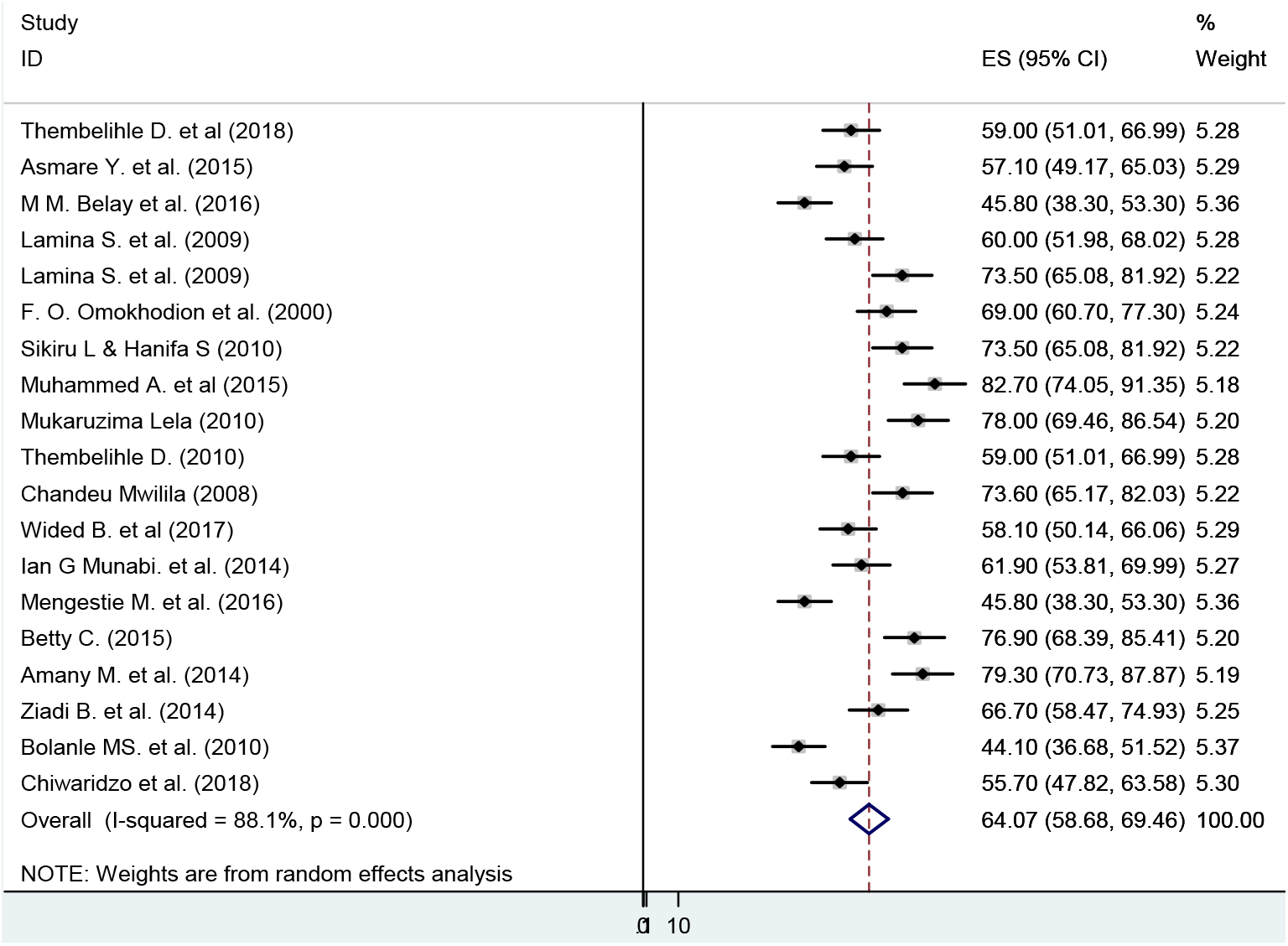
Forest plot of prevalence of low back pain among nurse in the African healthcare facilities, 2018.

### Subgroup analysis

According to the subgroup analyses, the highest prevalence of LBP among nurses was reported from west African region with prevalence rates of 68.46% (95% CI: 54.94–81.97; P-value <0.0001) and followed by north Africa region with prevalence rate of 67.95% (95% CI: 55.96–79.94; P value <0.0001) had the higher prevalence of LBP as compared to their south African counterparts, 59% (95% CI: 51–66.9; P-value <0.0001) (Figure 3).

**Figure 3:**
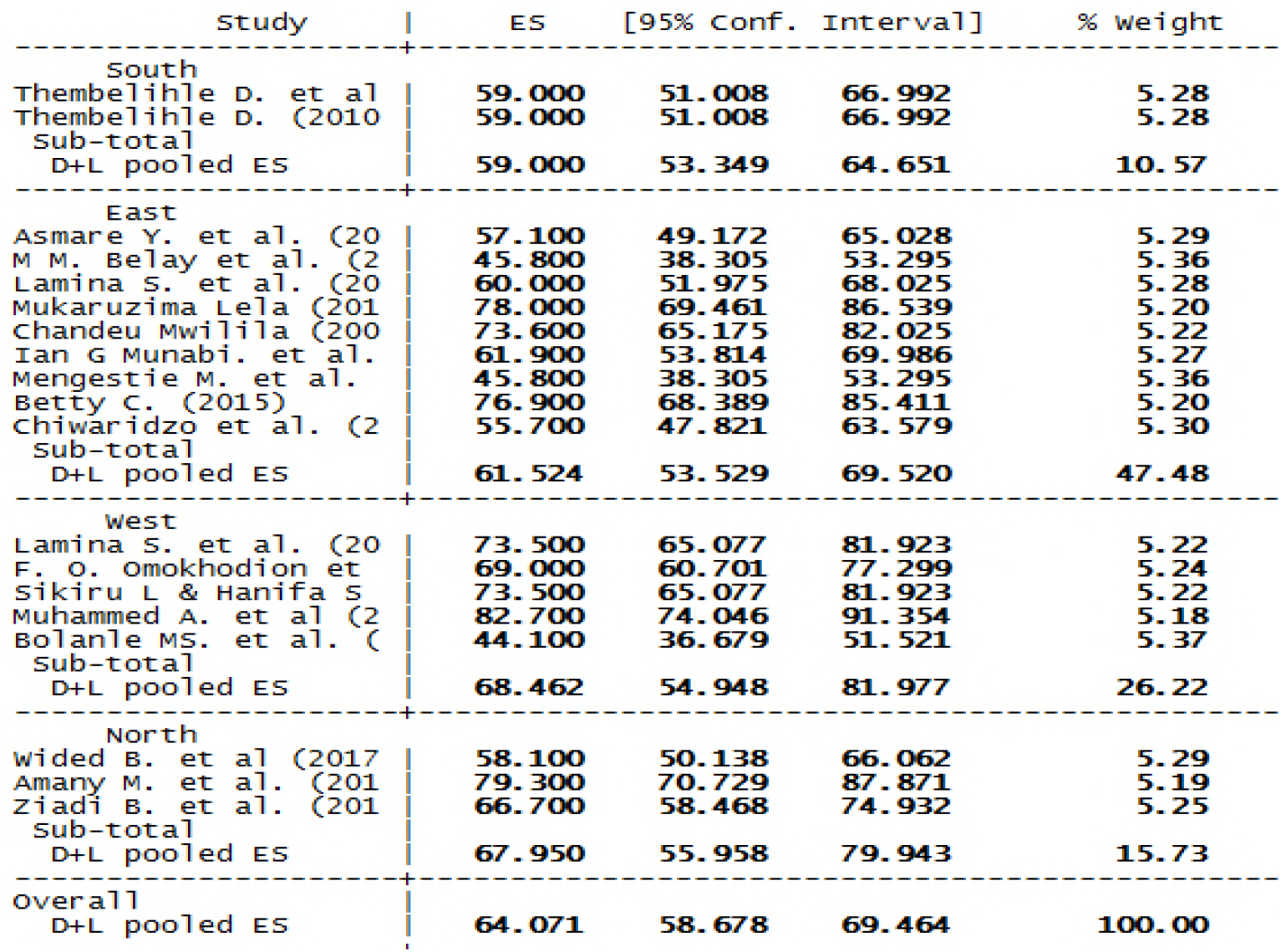
Subgroup analysis of low back pain among nurse using region of the continent in the African healthcare facilities, 2018.

### Meta–regression

Meta-regression analysis showed that there was no significant statistical relationship between the year of publication and the prevalence of the LBP (β = −0.82, P-value = 0.808) (Figure 4).

**Figure 4:**
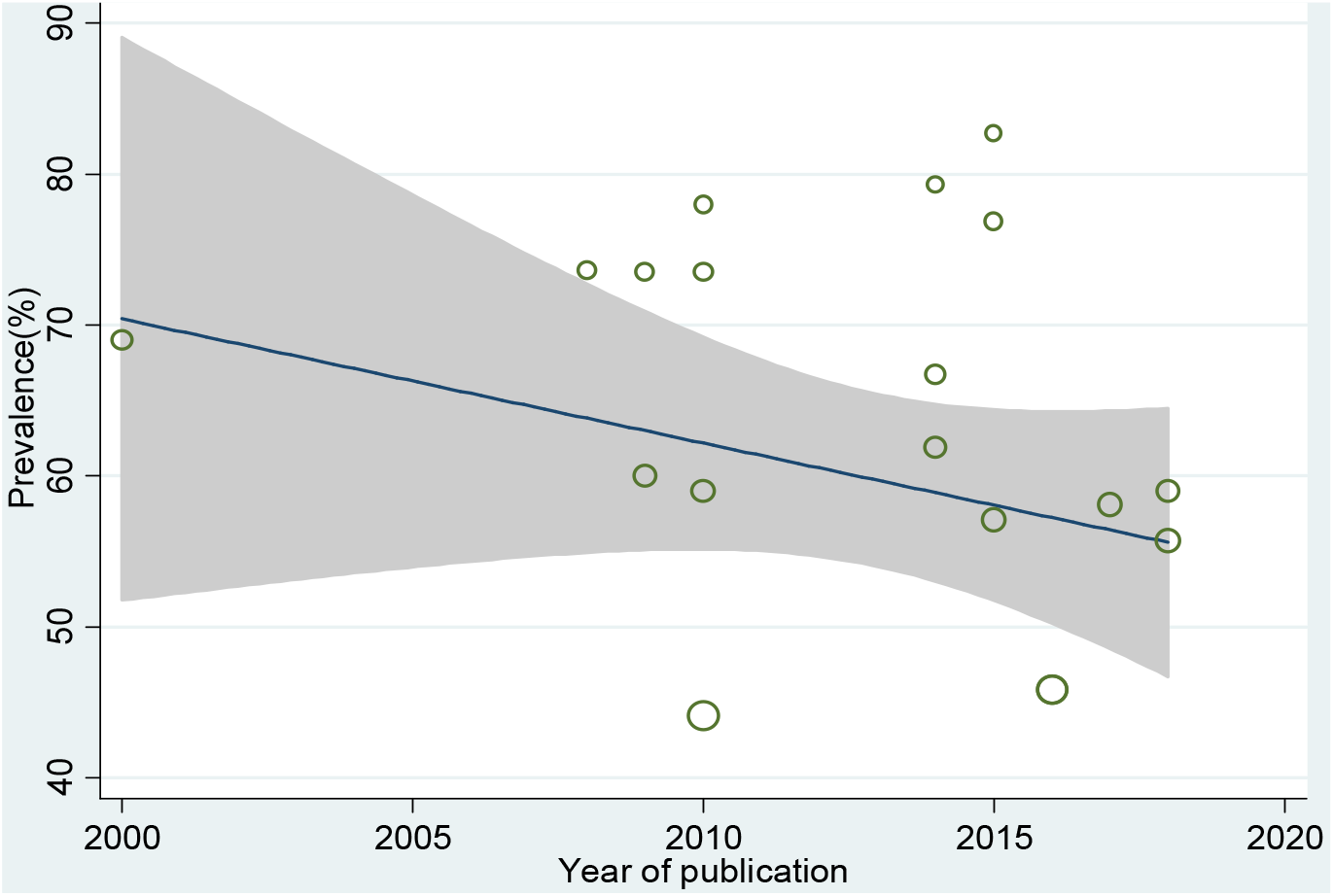
Funnel plot showing the relation between year of publication and prevalence of LBP among nurses working in different African health facilities, 2018.

The meta-regression also showed that there was no significant statistical relationship between the sample size and the prevalence of the LBP (β = −0.007, P-value = 0.93) (Figure 5).

**Figure 5:**
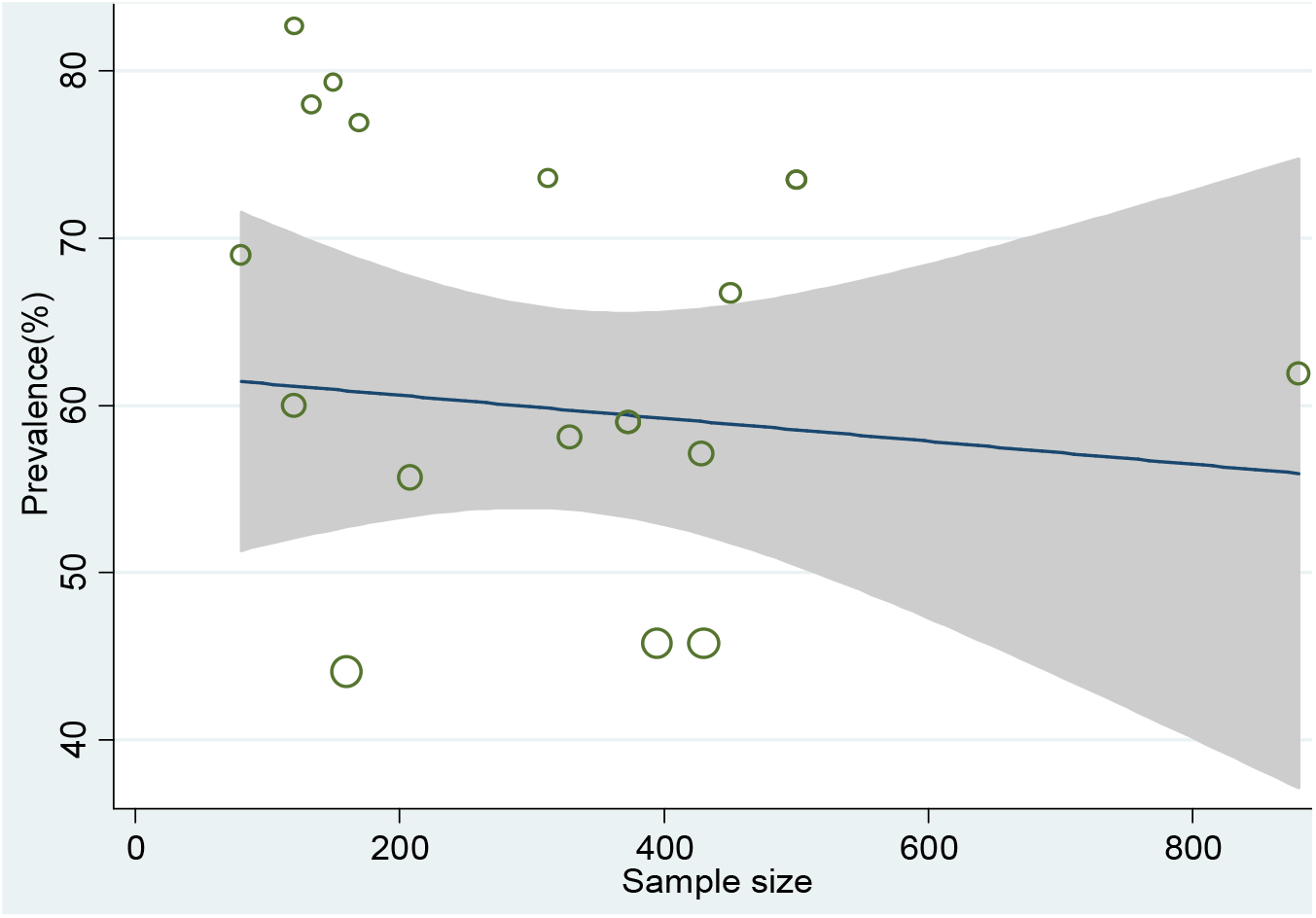
Funnel plot showing the relation between sample size and prevalence of LBP among nurses working in different African health facilities, 2018.

To assess publication bias, the funnel plot and the Egger’s test was conducted in the metaanalysis. The funnel plot and Egger’s regression tests (β = −0.0024, SE=0.06, P=0.96) showed that no evidence of publication bias for the included studies (Figure 6).

**Figure 6:**
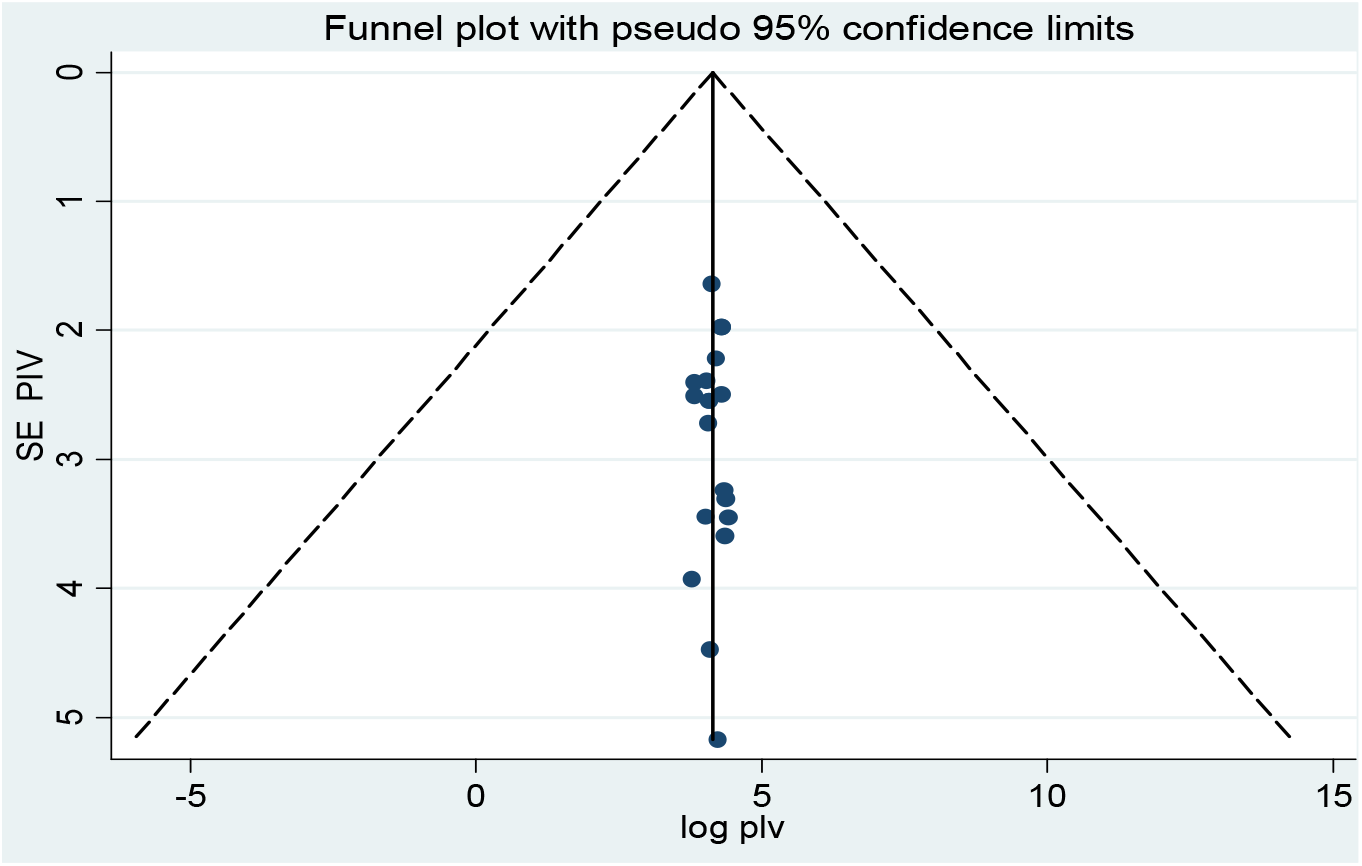
Funnel plot for assessing publication bias among studies, 2018

## Discussion

Low back pain is a common work-related musculoskeletal disorders in healthcare workers, with particularly imposing high risk on nursing professionals across different healthcare facilities mainly in hospital settings (68,69). Such problems are reported to considerably influence on quality of life of healthcare professionals and this will in turn affects the healthcare quality (29).

This study denotes the first effort to report on the prevalence of LBP among nurses working in healthcare facilities in the African continent. The aim of this systematic review and metaanalysis was to determine the pooled prevalence of LBP among nurse working at different healthcare facilities in different African regions. By providing a comprehensive picture, this study would help to recognize the impact of the problem on nurse professionals in African countries.

Low Back Pain is a regular occupational problem for nurses worldwide, and has been previously reported at rates between 45% in England (70), 63% in Australia (71). Research from Hong Kong and China has also showed that LBP may affect between 40.6% (72) and 56% (73) respectively. African studies report LBP rates between 44.1%, 79.4% and 82.7% (28,43,57,74)

The result of the present systematic review and meta-analysis carried on professional nurses working at different regions of African healthcare facilities showed that the overall prevalence of LBP among nurses was 64.07%. This finding was higher than a study done in Zimbabwe (67), Nigeria (57), Tunisia (10) and Iran (49) showed that the overall prevalence of LBP among nurses was 55.67%, 44.1%, 51.1% and 61.2% respectively.

Whereas a significant amount of studies conducted in the western nations and Asian countries revealed that the overall prevalence of LBP among nurses was higher when we compared to the present finding. A study done in Japan (75), Turkey (13) and United States of America (76) showed that the overall prevalence of LBP among nurses was 91.9%, 77.1% and 72.5% respectively which all indicated that the existence of more sever prevalence of LBP compared to the present study. Another study done in Swiss (77) and Italy (78) also revealed that the overall annual prevalence of LBP among nurses was found to be 73-76% and 86% respectively. This also confirms that there is higher prevalence of LBP among nurse in the western nations.

Studies done in the western and in some Asian countries revealed that the existence of higher prevalence of LBP among nurses. This finding is confirmed by different literatures in the subject area. This high existence of LBP among nurses in the developing nations might be high workloads (79) for patient care, conducting advanced procedures in different advanced area of patient care and this all might lead them to the experiencing of LBP in their working environment.

The results of this study identified the presence of high prevalence of low back pain among nurses who were working in the western region of Africa. In the present study, five different studies were incorporated from the western region of Africa and all of them were from Nigeria. Nigeria is the number one populous country in Africa. This has its own impact in the healthcare system including to the healthcare professionals. As it was mentioned in many literatures, nurses are the number one frontline healthcare professionals that can contact clients. This would have its own share to the development of LBP on nursing staffs.

### Limitation

Adequate studies were not incorporated from some region of the continent and even most of the studies were concentrated in a single country in each region of the continent. This might have its own shortfalls in producing the overall picture of the problem to the continent as a whole.

### Conclusion

Even though the overall prevalence of the present study is lower when compared to the western and Asian studies, it indicated that the prevalence of low back pain among nurses is on the move.

Conducting a study at national levels in order to determine the physical, mental, supervisor support, nurse colleague interaction, psychological, and work setting stressors in the work environment of nurses and their relationship with low back pain should investigated. This would enhance on how to identify the risk factors and to design a detailed plans for the prevention and control of low back pain. All the efforts made would improve nurses’ sense of belongingness, retention, quality of patient care and even organizational culture.

## Declarations

### Availability of data and material

All data generated or analyzed during this study are included in this manuscript.

### Authors’ Contributions

AS was involved in the design of the study, data analysis, and interpretation of the findings, report writing, and paper preparation. YW & EA involved in the analysis and interpretation of the data, and review of the report. WA approved the manuscript and all authors read and approved the final paper. All authors contributed equally to this work

## Appendix

The critical appraisal tool utilized to assess the quality of the included studies.

|  |  |
| --- | --- |
| Part I | Is the final sample representative of the target population? |
|  | 1. At least one of the following must apply in the study: an entire target population, randomly selected sample, or sample stated to represent the target population. |
|  | 2. At least one of the following: reasons for non-response described, non responders described, comparison of responders and non-responders, or comparison of sample and target population |
|  | 3. Response rate and, if applicable, drop-out rate reported |
| Part II | Quality of the data? |
|  | 4. Were the data primary data of low back pain or was it taken from a survey not specifically designed for that purpose? |
|  | 5. Were the data collected from each adult directly or were they collected from a proxy? |
|  | 6. Was the same mode of data collection used for all subjects? |
|  | 7. At least one of the following in case of questionnaire: a validated questionnaire or at least tested for reproducibility. |
|  | 8. At least one of the following in the case of an interview: Interview validated, tested for reproducibility, or adequately described and standardized. |
|  | 9. At least one of the following in the case of an examination: Examination validated, tested for reproducibility, or adequately described and standardized. |
| Part III | Definition of low back pain (LBP) |
|  | 10. Was there a precise anatomic delineation of the lumbar area or reference to an easily obtainable article that contains such specification? |
|  | 11. Was there further useful specification of the definition of LBP, or question(s) put to study subjects quoted such as the frequency, duration or intensity, and character of the pain. Or was there reference to an easily obtainable article that contains such specification? |
|  | 12. Were recall periods clearly stated: e.g., 1 week, 1 month or lifetime? |

